# Motor unit tracking using blind source separation filters and waveform cross-correlations: reliability under physiological and pharmacological conditions

**DOI:** 10.1101/2023.04.25.538344

**Authors:** Benjamin I. Goodlich, Alessandro Del Vecchio, Justin J. Kavanagh

## Abstract

Recent advancements in the analysis of high-density surface electromyography (HDsEMG) have enabled the identification, and tracking, of motor units (MUs) to study muscle activation. This study aimed to evaluate the reliability of MU tracking using two common methods: blind source separation filters and two-dimensional waveform cross-correlation. An experiment design was developed to assess physiological reliability, and reliability for a drug intervention known to reduce the firing rate of motoneurones (cyproheptadine). HDsEMG signals were recorded from tibialis anterior during isometric dorsiflexions to 10%, 30%, 50% and 70% of maximal voluntary contraction. MUs were matched within session (2 hr) using the filter method, and between sessions (7 days) via the waveform method. Both tracking methods demonstrated similar reliability during physiological conditions (e.g., MU discharge: filter ICC 10% of MVC = 0.76, to 70% of MVC = 0.86; waveform ICC: 10% of MVC = 0.78, to 70% of MVC = 0.91). Although reliability slightly reduced after the pharmacological intervention, there were no discernible differences in tracking performance (e.g., MU disc filter ICC: 10% of MVC = 0.73, to 70% of MVC = 0.75; DR waveform ICC: 10% of MVC = 0.84, to 70% of MVC = 0.85). The poorest reliability typically occurred at higher contraction intensities, which aligned with the greatest variability in MU characteristics. This study confirms that tracking method may not impact the interpretation of MU data, provided that an appropriate experiment design is employed. However, caution should be used when tracking MUs during higher intensity isometric contractions.

**NEW AND NOTEWORTHY:** The most direct way to validate longitudinal tracking of motor unit data extracted from high-density surface electromyography is to contrast findings with intramuscular electromyography. We use pharmacology to changes motor unit discharge properties as a non-invasive alternative to validate the reliability tracking motor units. This study confirmed that the specific tracking method may not impact interpretation of motor unit data at lower contraction intensities, however caution should be used when tracking units during higher intensities.

## INTRODUCTION

Recent developments in the analysis of high-density surface electromyograms (HDsEMG) have unlocked the potential to reveal how large populations of motor units regulate their recruitment and discharge during the performance of goal-directed tasks (1–5). However, a technical challenge exists for longitudinal studies where experiment designs often require multiple recording sessions to examine the effect of an intervention. Without appropriate motor unit tracking it is conceivable that different motor units will be assessed pre- and post-intervention. This presents a barrier for revealing how interventions influence the muscle activation and could hinder, rather than advance, our understanding of how the nervous system controls movement. Although precise mapping of muscle activation patterns has the potential to study interventions across a wide range of motor activities and neuromuscular disorders, the reliability of tracking motor units across time remains largely unknown.

There is now convincing evidence that the same motor units extracted from HDsEMG of the same muscle can be reliably identified in a single testing session when muscle contractions of different intensity are performed. This has been validated with the two-source method, so that the same motor unit is identified from intramuscular EMG signals (the current gold standard) as well as by blind source separation of the HDsEMG (6, 7). This validation is beneficial because it not only confirms the utility of within session tracking using blind source separation filters, but it also lays a platform to compare other methods to the blind source separation. A second popular method of motor unit tracking involves waveform correlation, where it is suggested that the high correlation in two-dimensional waveform across different experimental days is due to the tracking of the same motor unit (5). This has been demonstrated over a period of weeks and months (1, 5, 8). However, validating this technique with respect to the gold-standard method of intramuscular EMG is difficult, owing to its invasive nature and the logistical challenge of placing fine wire electrodes in precisely the same position for each testing session.

The identification of single motor-unit activity in surface EMG recordings is reliant on the unique representation of motor units by their surface action potentials, where invariant patterns of surface action potentials is believed to reflect the discharge of the same motor unit over time (9). However, interventions that alter muscle force, almost by definition, are designed to introduce variations in motor unit discharge properties, and often impact the morphology, temporal, and frequency characteristics of the motor unit action potential waveforms (MUAPs). For example, motor unit discharge is known to increase by ∼0.5 Hz with repeated muscle stretching (10), reduce by ∼0.3 Hz with stimulation induced reciprocal inhibition (11), reduce by ∼0.6 Hz with vibration induced reciprocal inhibition (12), and a four-week resistance training program is known to increase motor unit discharge by ∼3 Hz (1). Thus, it is important for any tracking method to be reliable when motor unit activity is predicted to change with an intervention. We have recently used pharmacological interventions to induce serotonergic receptor blockade, where antagonism of 5-HT_2_ receptors in the motor pathway reduce the discharge of tibialis anterior motor units by >1 Hz during both rapid (13) and steady state (14) isometric contractions. The observed suppression in discharge during steady state muscle activation was found to align with a decrease in of estimates of persistent inward current activity, indicative of a reduction in intrinsic motoneuron excitability. We use this knowledge of 5-HT_2_ antagonism in the current study to evoke a pharmacological-based reduction in motor unit firing frequency, so that the reliability of motor unit tracking can be determined when intrinsic mechanisms of motoneurone excitability are manipulated.

The purpose of this study was to evaluate the reliability of two common tracking procedures: the blind source separation filter method and the two-dimensional waveform correlation method. HDsEMG signals were recorded from the tibialis anterior during the performance of isometric dorsiflexions to four different submaximal intensities. The HDsEMG signal was decomposed via convolution kernel compensation (15) before each tracking method was applied to the same data set. The design of this study also included a pharmacological intervention with known effects on motor unit discharge characteristics so that the effect of exogenous manipulation of motoneurone excitability on tracking performance could be investigated. Given that there is unlikely to be large changes in motor units discharge characteristics between the tracked data sets (for both the physiological condition and the pharmacological intervention), we hypothesised that both techniques will demonstrate similar tracking reliability. The effect of 5-HT_2_ receptor blockade on motor unit discharge, motor unit recruitment, and motor unit decruitment, for the contraction protocol employed in this study has been previously reported (14).

## METHODS

### Participants and ethical approval

Eleven healthy, recreationally active individuals (age 24.1 ± 2.6 years, 4 female) were recruited to the study. Approval for testing procedures were obtained via Griffith University’s Human Research Ethics committee (GU Ref No: 2020/264), and all procedures were performed in accordance with the *Declaration of Helsinki*. Written informed consent was obtained for all participants prior to testing. Participants were screened for acute or chronic neuromuscular injury, as well as contraindications associated with cyproheptadine administration. Participants were instructed to refrain from any depressants or stimulants such as alcohol, caffeine, or moderate-to-high intensity exercise on the morning of testing.

### Experiment setup

Participants sat in a therapy chair which was adjusted for each participant to position the right hip, knee, and ankle at 90° of flexion in the sagittal plane. The participants’ right foot was secured with a non-compliant, ratchet type binding to a custom designed foot plate which incorporated a torque sensor (capacity = 565 Nm, Model 2110-5K; Honeywell International Inc., Charlotte, NC, USA). The foot plate was attached to a bespoke aluminium frame which was secured to the chair, ensuring that the torque sensor axis of rotation aligned to participants’ malleoli (Figure 1A). Ankle torque was sampled at 2000 Hz using a Power 1401 interface with Spike2 software (version 7, Cambridge Electronic Design Ltd., UK). Feedback for the unfiltered torque signal was displayed on a computer monitor positioned approximately 1 m in front of the participant, with dorsiflexion torque presented as a positive inflection in torque on the screen.

**Figure 1.**
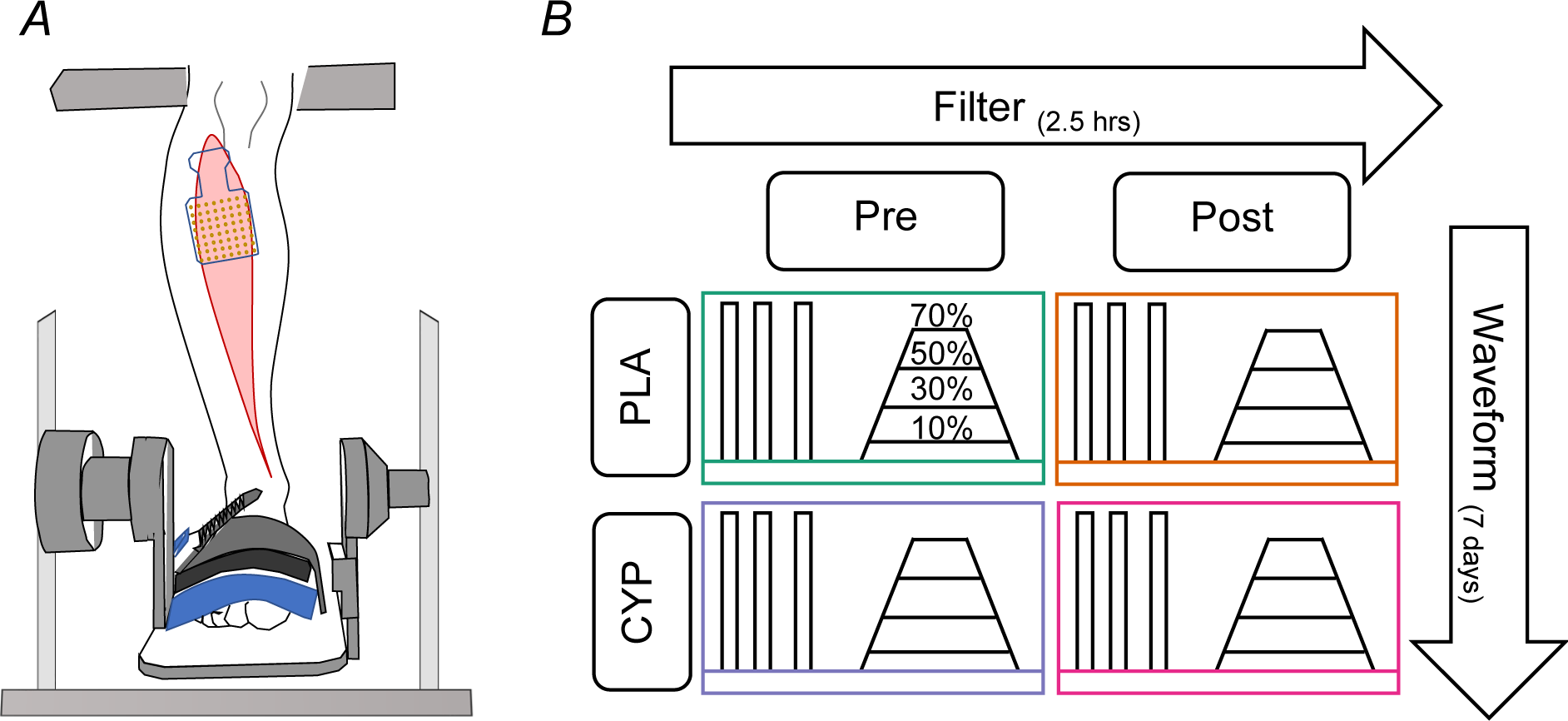
Experiment setup and contraction protocol. *A*, The right foot was secured to a foot plate attached to a torque sensor. HDsEMG electrodes were fixed over the TA muscle belly and oriented in the estimated direction of muscle fibres. *B,* Maximal and submaximal dorsiflexions were performed at four time points: pre-placebo (green), post-placebo (orange), pre-cyproheptadine (purple) and post-cyproheptadine (pink). Submaximal contractions were performed to 10%, 30%, 50% and 70% of MVC; the order of which was randomised. Submaximal contraction durations were 24 s for all intensities. The ramp up and down rate of the trapezoidal contractions was fixed at 10% of MVC/s. Motor units were matched intra-session across the 2.5 hr pharmacological intervention using blind source separation decomposition techniques (filter method). Motor units were matched inter-session across 7 days using 2D waveform cross-correlation (waveform method). HDsEMG, high-density surface electromyography; TA, tibialis anterior; MVC, maximal voluntary contraction; PLA, placebo; CYP, cyproheptadine.

Muscle activity for the tibialis anterior was measured using a semi-disposable 64-channel (8 퀇 8) HDsEMG grid electrode with a 10 mm inter-electrode distance (OTBioelettronica, Torino, Italy). Following skin preparation (shaving, abrasion, and cleansing with 70% isopropyl alcohol), the position and orientation of the electrode grid was determined by an experienced investigator via palpation of the right tibialis anterior muscle belly (Figure 1A). Electrodes were fixed to the middle of the muscle belly using a bi-adhesive, perforated foam layer and conductive paste (SpesMedica, Battipaglia, Italy). A dampened strap ground electrode (OTBioelettronica, Torino, Italy) was positioned over the right ankle malleoli. HDsEMG signals were recorded in monopolar mode and converted to digital signal by a 16-bit wireless amplifier (Sessantaquattro, OTBioelettronica, Torino, Italy). HDsEMG signals were recorded and visualised using OTBioLab+ software (version 1.5.5., OTBioelettronica, Torino, Italy).

### Experiment protocol

Participants attended two testing sessions separated by one week. Figure 1B illustrates the contraction protocols that were used in each session, whereby TA EMG data could be examined using two motor unit tracking techniques. The contraction protocols remained consistent throughout all data collection. Each participant performed 5 maximal effort dorsiflexions for ∼3 s (with ∼ 3 min rest between contractions) to establish maximal voluntary contraction (MVC) torque. The trial that generated the highest magnitude of torque was determined to be the participant’s MVC. Participants then performed trapezoidal contractions to 10%, 30%, 50% and 70% of this MVC. Each trapezoid was 24 s in duration, where rate of torque increase and rate of torque decrease was 10% MVC/s. Contraction duration and rate of torque development were fixed to keep motor unit spike frequency adaptation and spike threshold accommodation consistent across intensities (16–18). The trapezoidal contractions were presented on a monitor placed 1 m in front of the participant, where trials were only accepted if the participant’s dorsiflexion torque remained within 1% of the prescribed trajectories.

In one session the series of muscle contractions were performed pre- and post-ingestion of a placebo, and in the other session the same series of contractions were performed pre-and post-ingestion of a serotonin receptor blocking drug. Depending on the session, a single oral dose of placebo (containing Avicel filler) or a single dose of cyproheptadine (8 mg) was administered. Post-pill torque and EMG measurements were made 2.5 hr after oral administration. The placebo and cyproheptadine were compounded in opaque capsules to ensure blinding of the drug condition. The timing of testing aligned with high plasma concentrations of cyproheptadine (19, 20), as well as the testing window reported in previous cyproheptadine studies (13, 14, 21). The order that participants were allocated to the placebo or drug session was counterbalanced.

### Motor unit decomposition

Monopolar HDsEMG signals were digitally band pass filtered at 20–500 Hz with a second-order Butterworth filter. HDsEMG signals were then decomposed into individual motor unit action potentials using blind source separation, via the convolutive kernel compensation method (15). This method has been validated previously for a broad range of contraction intensities of the tibialis anterior muscle (1, 3, 7, 13, 22). The decomposition accuracy was assessed using pulse-to-noise ratio dB during each individual contraction (7), and decomposed spike trains showing pulse-to-noise ratios < 28 dB were discarded from the analysis, as described in Del Vecchio, Holobar (2). All motor unit pulse trains were manually inspected and only pulse trains with a reliable discharge pattern were considered for analysis. Motor unit recruitment threshold and derecruitment threshold was calculated as the torque value corresponding to the first and last motor unit firing, respectively. Motor unit discharge rate was calculated during the plateau phase of the trapezoidal contraction (the average discharge rate of the first 10 interspike intervals during steady contraction at the target intensity).

### Tracking motor unit activity with motor unit decomposition filters

Intra-session tracking was achieved with the abovementioned blind source separation, whereby motor units were tracked from **pre- to post-pill** ingestion for the same contraction intensity and drug condition (Figure 1B). This tracking routine was performed for both the placebo and cyproheptadine testing sessions. This was achieved by concatenating the HDsEMG recordings from pre- and post-ingestion, and then applying the individual motor unit filters to the entire concatenated data set. Intra-session tracking will henceforth be referred to as **filter tracking**.

### Tracking motor unit activity with waveform correlation analysis

Motor unit action potentials extracted during EMG decomposition were matched longitudinally from session one to session two via two dimensional cross-correlation of motor unit action potential waveforms (1, 5). Motor unit action potential waveforms were estimated with spike triggered averaging, and the spatial representation of the action potential waveforms were cross correlated between ramp contractions. Longitudinal inter-session tracking was performed within drug condition and contraction intensity. Specifically, null comparisons were made between motor units tracked from **pre-placebo to pre-cyproheptadine**, and drug comparisons were made between motor units tracked from **post-placebo to post-cyproheptadine** (Figure 1B). Inter-session tracking will henceforth be referred to as **waveform tracking**.

### Random matching of motor units for filter tracking and waveform tracking

As a comparator to the accepted methods of matching motor units acutely and longitudinally, motor units were also matched randomly. Random matching was achieved using bespoke code (MATLAB (R2020a), The Mathworks Inc., Natick, MA) which randomly paired the decomposed motor units from each subject within drug condition and contraction intensity. Random matching was performed to assess the reliability of comparing motor unit properties in both placebo and drug conditions in the absence of a specific method of ensuring the same motor unit are tracked across the comparison. We have then compared the physiological properties of the motor units before and after placebo or cyproheptadine, with the expectation being that the use of random matching will be far less reliable than the two accepted methods of matching.

### Statistical analysis

All statistical analysis was performed in R, using RStudio (version 4.1.1; R Foundation for Statistical Computing, Vienna, Austria). The motor unit characteristics assessed were recruitment threshold, average discharge rate during the plateau, and derecruitment threshold. These variables were analysed at each contraction intensity (10%, 30%, 50% and 70% of MVC) for both the filter and waveform tracking methods. To assess test-retest reliability of motor unit characteristics for each tracking method, two-way mixed effects, consistency, single measurement intraclass correlation coefficients (ICC 3,1) were computed. Additionally, within subject variability was calculated using the mean intra-participant coefficient of variation 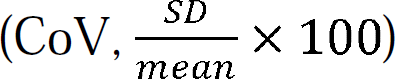 and the standard error of measurement 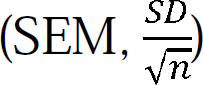. Change scores were calculated for each motor unit property as matched using both tracking methods. To visualise this information and describe the agreement between the two tracking methods, separate Bland-Altman plots were generated for each pharmacological state (placebo and cyproheptadine). Specifically, data for each motor unit is plotted on the ordinate as the difference in change score between the filter and tracking methods, and on the abscissa as the average change score between the filter and waveform methods. To assess the strength of bivariate correlations between motor units matched across conditions using filter, waveform, and random tracking methods, a Pearson product-moment correlation was used. This analysis was performed for all motor unit properties at each contraction intensity during both pharmacological states.

## RESULTS

### Motor unit decomposition

A total of 1052 motor units were identified via decomposition of HDsEMG signals from the submaximal dorsiflexions. Using the filter method, 296 units were successfully matched from pre- to post-placebo and 344 units matched pre- to post-during the pharmacological intervention. Using the waveform method, 196 units were successfully matched from pre- to post-placebo and 216 units matched pre- to post-during the pharmacological intervention. For the placebo session, the number of motor units identified per subject ranged from 6-20 at 10% MVC, 3-20 at 30% MVC, 3-13 at 50% MVC, and 2-9 at 70% MVC. For the cyproheptadine session, the number of motor units identified per subject ranged from 3-19 at 10% MVC, 2-21 at 30% MVC, 2-12 at 50% MVC, and 2-8 at 70% MVC.

Data for each decomposed motor unit are presented in Figure 2. Motor unit discharge rate, recruitment threshold, and derecruitment threshold, all increase with increasing contraction intensity irrespective of the drug condition. There is also a greater range of physiological values at the higher contraction intensities for each motor unit property. The pharmacological intervention caused a significant reduction in motor unit discharge rate and a significant increase in derecruitment threshold, which can be observed in Figure 3 as a leftward shift of the discharge rate change scores and a subtle rightward shift of the derecruitment threshold change scores.

**Figure 2.**
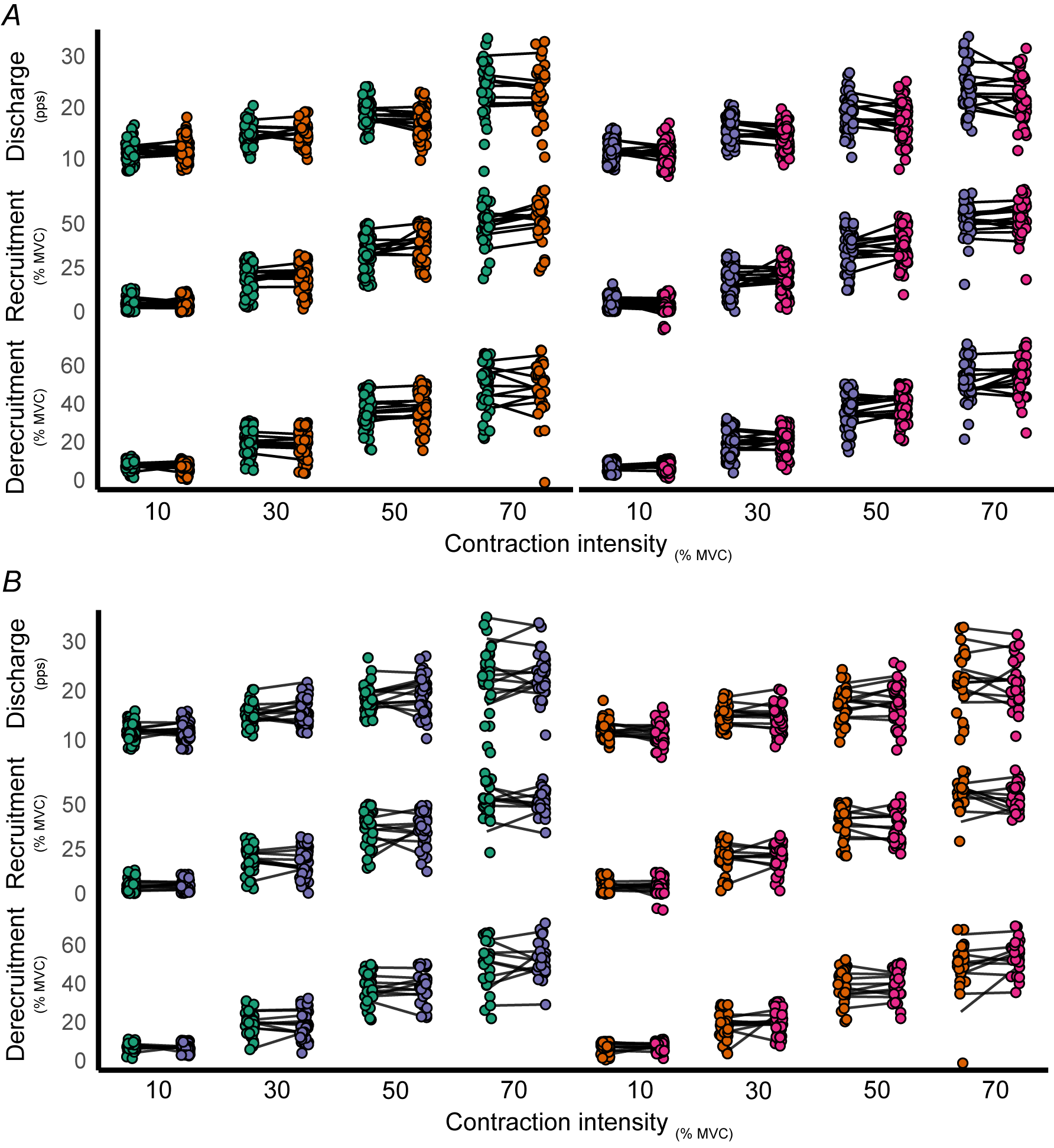
Motor unit discharge rate, recruitment threshold, and derecruitment threshold. Motor roperties were extracted from tibialis anterior HDsEMG at four timepoints: pre-placebo (green), placebo (orange), pre-cyproheptadine (purple) and post-cyproheptadine (pink). Motor units were hed over a 2.5 hr period using the filter method from pre- to post-placebo and pre- to post-heptadine, shown in *A* (*n* = 11 subjects). Motor units were matched over 7 days using the form method from pre-placebo to pre-cyproheptadine and post-placebo to post-cyproheptadine, n in *B* (*n* = 11 subjects). Each point represents an individual motor unit, the colour of which sents the timepoint at which data were collected. The solid black lines represent subject averages ch timepoint.

**Figure 3.**
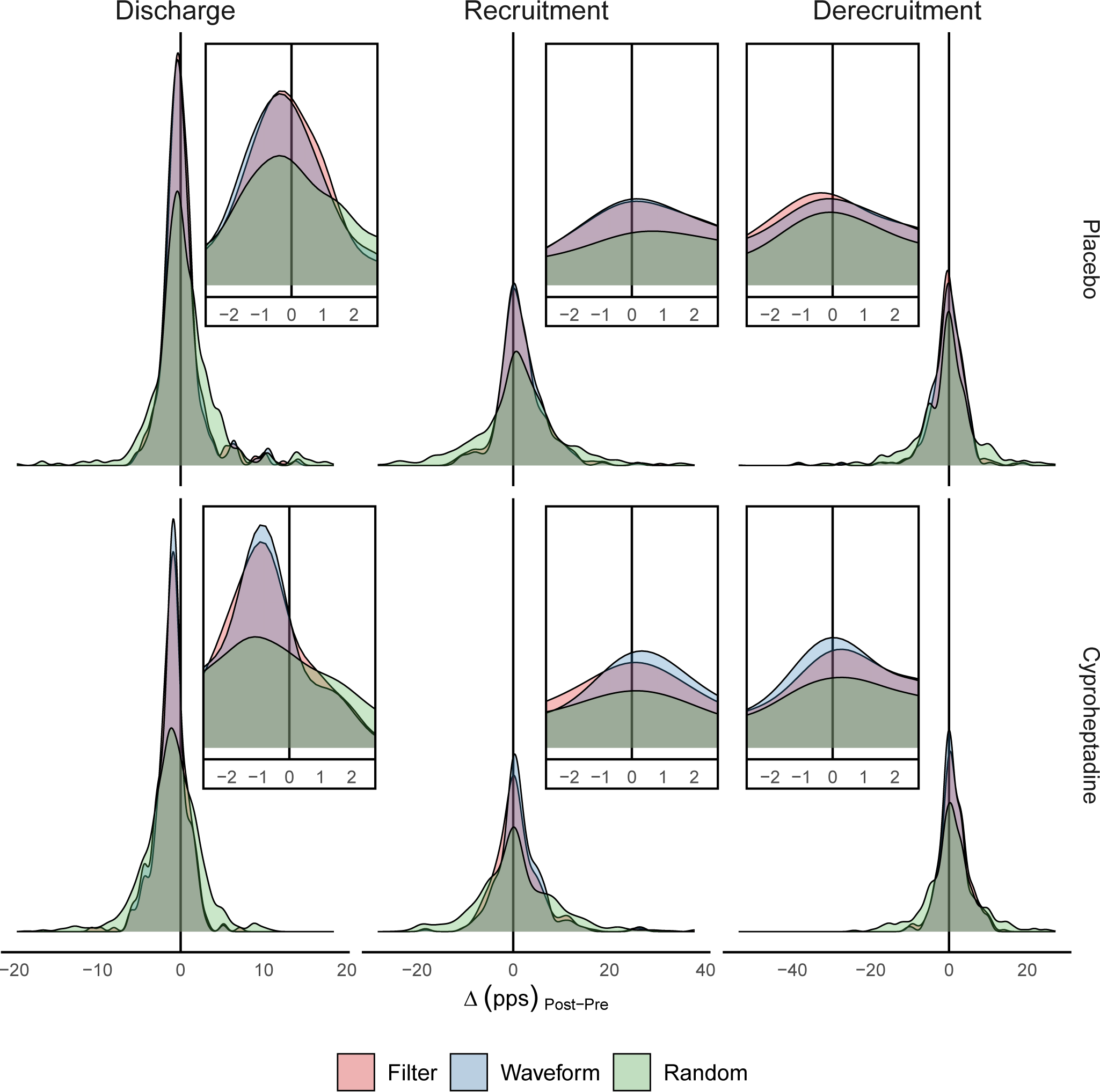
Change score density distributions. Density plots represent the distribution of changes s for motor units tracked from pre to post pill ingestion. Change scores are pooled for all action intensities (10%, 30%, 50% and 70% of MVC). Rows represent the drug condition and mns represent different motor unit properties (discharge rate in pulses per second, recruitment hold and derecruitment threshold in % of MVC). Change score distributions for the filter method hown in red, the waveform method is shown in blue, and the randomly matched condition is n in green. The insert shows the peaks of each distribution for each property and pharmacological Note the leftward shift present in the discharge rate change scores during the cyproheptadine ition.

### Reliability of filter tracking and waveform tracking method: placebo

Data obtained in the placebo session represent normal physiological responses during voluntary dorsiflexions. The filter tracking method was used to assess inter-session reliability, and the waveform method was used to assess intra-session reliability of the same motor units. Table 1 describes the reliability, error, and variability of motor unit discharge rate, recruitment and derecruitment threshold for units matched across the placebo condition using both the filter and waveform tracking methods. In general, both the ICC and SEM increased with increasing contraction intensity for each of the motor unit properties as matched using both methods. The CoV was relatively consistent for discharge rate at 10%, 30% and 50% of MVC, however increased during 70% of MVC when matched using both methods.

**Table 1.** Placebo intra- and intersession reliability of motor unit parameters.

| Variable | Intensity<br>(%MVC) | Filter |  |  | Waveform |  |  |
| --- | --- | --- | --- | --- | --- | --- | --- |
|  |  | ICC | SEM | CoV<br>(%) | ICC | SEM | CoV<br>(%) |
| Discharge rate<br>(pps) | 10 | 0.76 | 0.10 | 12.54 | 0.78 | 0.13 | 10.62 |
|  | 30 | 0.68 | 0.16 | 11.58 | 0.70 | 0.19 | 10.10 |
|  | 50 | 0.83 | 0.26 | 12.95 | 0.84 | 0.37 | 10.99 |
|  | 70 | 0.86 | 0.56 | 15.80 | 0.91 | 0.80 | 15.68 |
| Recruitment<br>(%MVC) | 10 | 0.51 | 0.20 | 76.10 | 0.74 | 0.28 | 81.14 |
|  | 30 | 0.87 | 0.60 | 33.31 | 0.85 | 0.76 | 31.00 |
|  | 50 | 0.72 | 0.89 | 23.62 | 0.79 | 1.17 | 22.21 |
|  | 70 | 0.91 | 1.16 | 17.63 | 0.86 | 1.21 | 12.96 |
| Derecruitment<br>(%MVC) | 10 | 0.36 | 0.14 | 28.99 | 0.21 | 0.20 | 32.23 |
|  | 30 | 0.87 | 0.53 | 32.76 | 0.83 | 0.58 | 27.12 |
|  | 50 | 0.93 | 0.77 | 19.05 | 0.94 | 1.00 | 14.29 |
|  | 70 | 0.82 | 1.37 | 21.51 | 0.80 | 1.80 | 18.89 |

ICC, intraclass correlation coefficient; SEM, standard error of measurement; CoV, coefficient of variation; pps, pulses per second; MVC, maximal voluntary contraction

Bland-Altman plots were constructed to investigate the level of agreement between the two tracking methods for measuring change in each motor unit property at the different submaximal intensities. Figure 4 illustrates the agreement of the two tracking methods for the placebo condition. At 10% of MVC, a high level of agreement is demonstrated for all motor unit properties by the clustering of datapoints around zero on both axes. This is to be expected in the placebo condition, where there should be minimal change in the behaviour of motor units. As the contraction intensity increases, the data points become more sparce, illustrating reduced agreement and a higher level of variability in the change scores measure across all motor unit properties assessed.

**Figure 4.**
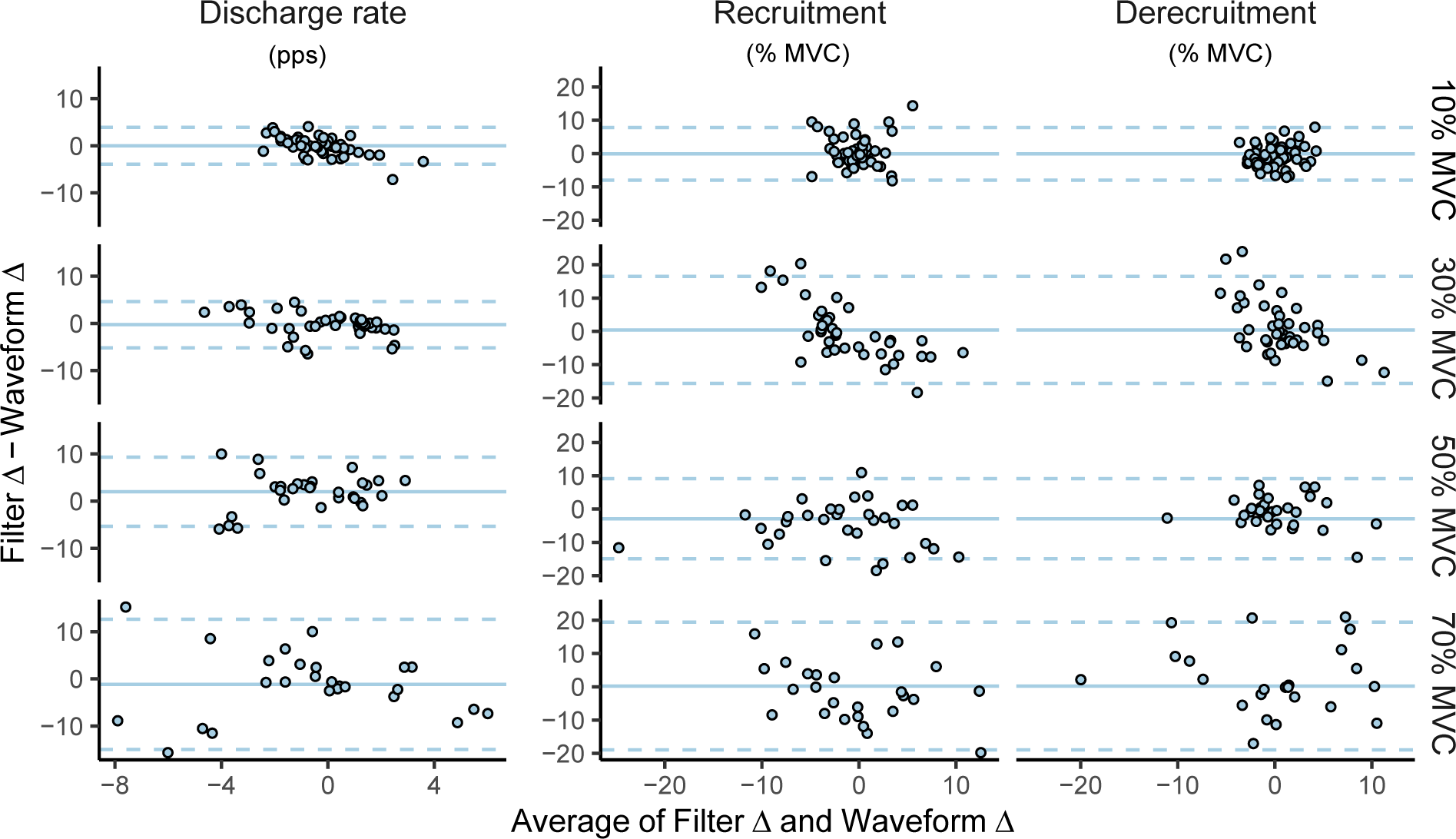
Bland-Altman plot for placebo data. Each point represents a change score of a motor unit h has been tracked across the placebo intervention using both the filter and waveform methods. for each motor unit is plotted on the ordinate as the difference in change score between the filter racking methods, and on the abscissa as the average change score between the filter and waveform ods. Columns represent discharge rate, recruitment threshold and derecruitment threshold, and represent the different contraction intensities (10%, 30%, 50% and 70% of MVC). The solid blue represent the average difference in change scores, and the dashed blue lines represent the 95% dence interval around the mean difference. Discharge rate data is presented as pulses per second, and recruitment/derecruitment data is presented relative to subjects’ maximal voluntary action (% MVC).

Figure 5 shows the subject-wise *r*^2^ values achieved by performing Pearson product-moment correlations between motor unit properties for the placebo condition. Data are presented for matching via the filter method, waveform method and random matching of motor units. Unsurprisingly, correlations between randomly matched units consistently returned low *r*^2^ values, irrespective of motor unit property being assessed or contraction intensity being performed. Correlations between units matched via the filter and waveform method demonstrated similarly high *r*^2^ values. However, *r*^2^ values tended to become more variable with increasing contraction intensity.

**Figure 5.**
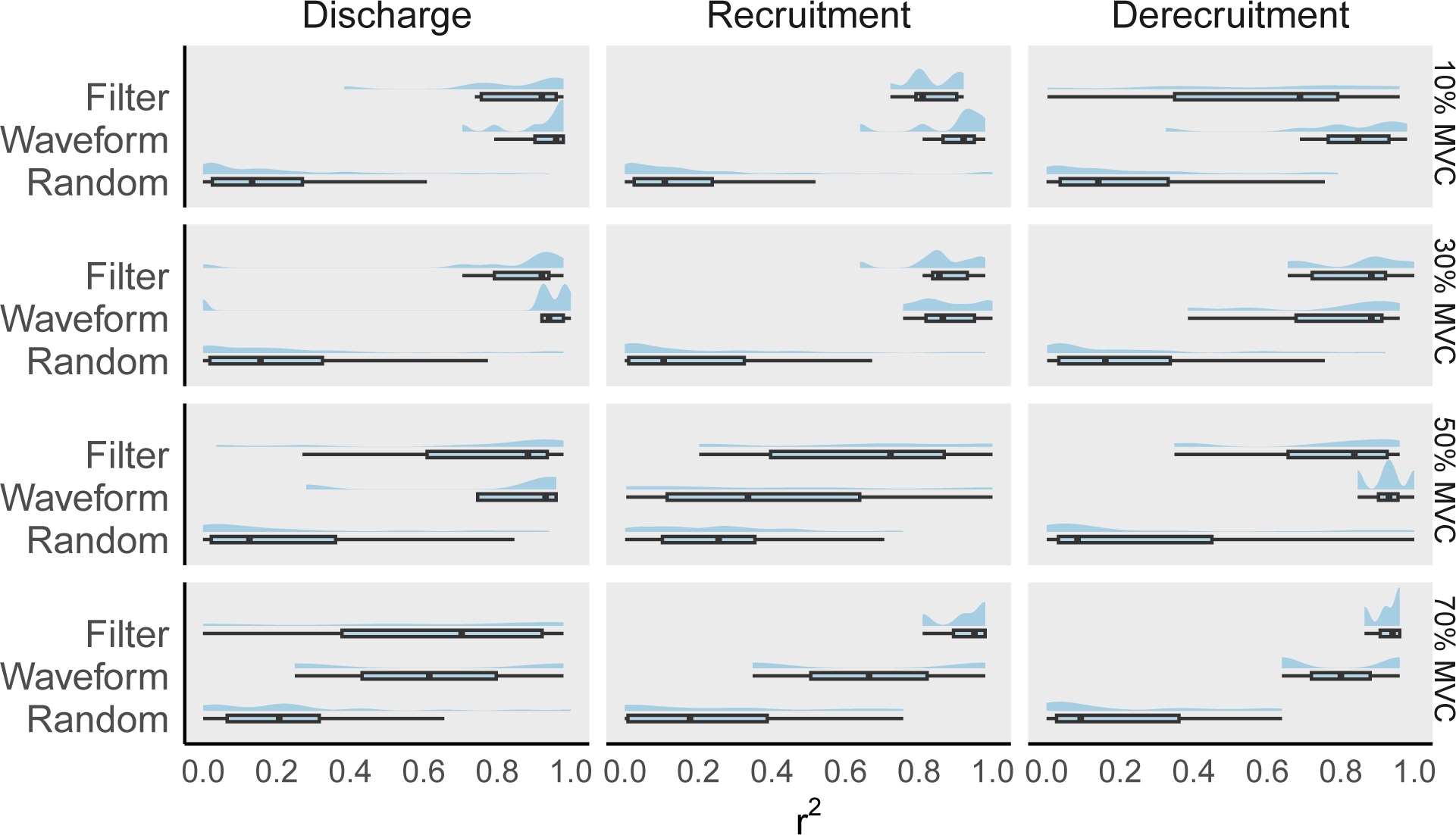
Pearson correlation coefficients for placebo data. Box plots and density plots represent the bution of individual subject correlation coefficients for motor units which has been tracked across lacebo intervention using the filter, waveform, and random methods. The distribution of r^2^ values played on the abscissa, and distributions are grouped on the ordinate by tracking method. Columns sent discharge rate, recruitment threshold and derecruitment threshold, and rows represent the ent contraction intensities (10%, 30%, 50% and 70% of MVC).

### Reliability of filter tracking and waveform tracking method: pharmacological intervention

Data obtained in the drug session represent physiological change due to serotonergic receptor blockade during voluntary dorsiflexions. The reliability, error, and variability of motor unit discharge rate, recruitment and derecruitment threshold for units matched across the cyproheptadine condition using both the filter and waveform tracking methods are presented in Table 2. Similar to the placebo condition, there was a general increase in the ICC and SEM with increasing contraction intensity for each of the motor unit properties as matched using both methods. The ICCs in the cyproheptadine condition were typically lower than the placebo condition for motor unit discharge rate and derecruitment threshold. The CoV was comparable between the placebo and cyproheptadine conditions, behaving similarly across the different motor unit properties and contraction intensities.

**Table 2.** Cyproheptadine intra- and intersession reliability of motor unit parameters.

| Variable | Intensity<br>(%MVC) | Filter |  |  | Waveform |  |  |
| --- | --- | --- | --- | --- | --- | --- | --- |
|  |  | ICC | SEM | CoV | ICC | SEM | CoV |
| Discharge rate<br>(pps) | 10 | 0.55 | 0.12 | 14.36 | 0.70 | 0.13 | 13.25 |
|  | 30 | 0.78 | 0.16 | 12.43 | 0.86 | 0.24 | 10.26 |
|  | 50 | 0.79 | 0.29 | 13.87 | 0.83 | 0.44 | 13.84 |
|  | 70 | 0.73 | 0.44 | 11.61 | 0.84 | 0.57 | 9.79 |
| Recruitment<br>(%MVC) | 10 | 0.69 | 0.25 | 86.61 | 0.72 | 0.30 | 96.31 |
|  | 30 | 0.81 | 0.54 | 34.99 | 0.75 | 0.73 | 30.17 |
|  | 50 | 0.74 | 0.88 | 23.97 | 0.83 | 1.11 | 19.38 |
|  | 70 | 0.82 | 0.92 | 12.64 | 0.83 | 0.95 | 8.83 |
| Derecruitment<br>(%MVC) | 10 | 0.54 | 0.15 | 30.58 | 0.64 | 0.18 | 30.30 |
|  | 30 | 0.84 | 0.45 | 27.95 | 0.83 | 0.65 | 28.04 |
|  | 50 | 0.89 | 0.78 | 20.74 | 0.87 | 0.98 | 14.79 |
|  | 70 | 0.75 | 0.92 | 12.17 | 0.85 | 1.16 | 7.69 |

ICC, intraclass correlation coefficient; SEM, standard error of measurement; CoV, coefficient of variation; pps, pulses per second; MVC, maximal voluntary contraction

Figure 6 illustrates the agreement between the two tracking methods for measuring change in each motor unit property at the different submaximal intensities for the drug condition. At 10% of MVC, a high level of agreement is demonstrated by the clustering of datapoints for each motor unit property. It is worth noting the leftward shift of the discharge rate data points and a subtle rightward shift of the derecruitment data points. Again, this is a product of the average changes to these properties with the pharmacological intervention. Similar to the placebo condition, an increase in contraction intensity sees the data points become more sparce, once again illustrating reduced agreement and a higher level of variability in the change scores measure across all motor unit properties assessed.

**Figure 6.**
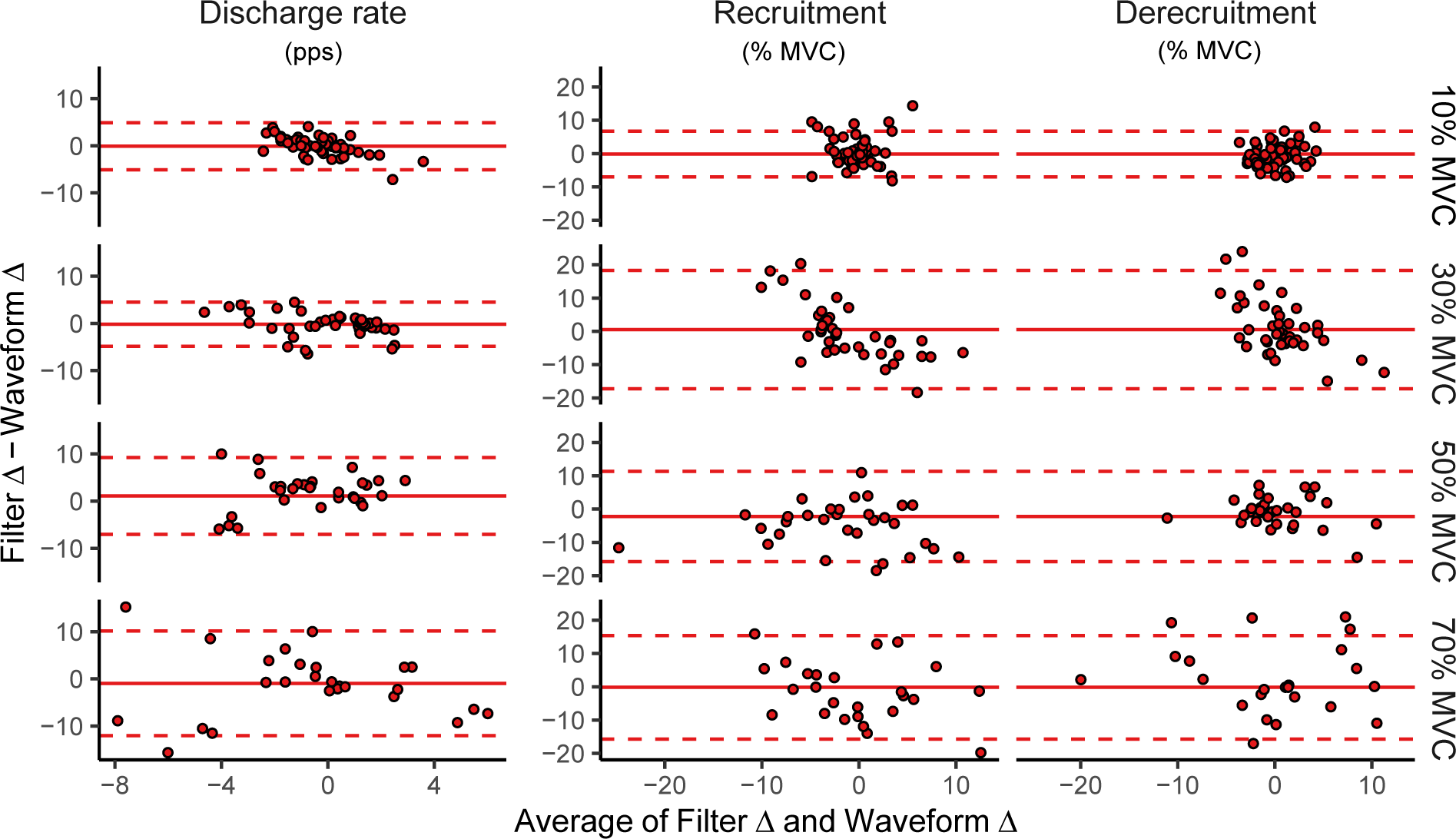
Bland-Altman plot for drug data. Each point represents a change score of a motor unit h has been tracked across the cyproheptadine intervention using both the filter and waveform ods. Data for each motor unit is plotted on the ordinate as the difference in change score between lter and tracking methods, and on the abscissa as the average change score between the filter and form methods. Columns represent discharge rate, recruitment threshold and derecruitment hold, and rows represent the different contraction intensities (10%, 30%, 50% and 70% of MVC). olid red lines represent the average difference in change scores, and the dashed red lines represent 5% confidence interval around the mean difference. Discharge rate data is presented as pulses per d (pps), and recruitment/derecruitment data is presented relative to subjects’ maximal voluntary action (% MVC).

The subject-wise *r*^2^ values achieved by performing Pearson product-moment correlations between motor unit properties before and after the cyproheptadine condition are presented in Figure 7. Once again, correlations between randomly matched units consistently returned low *r*^2^ values, irrespective of motor unit property being assessed or contraction intensity being performed. Units matched via the filter and waveform method demonstrated similarly high *r*^2^ values during the cyproheptadine condition, however, the same trend of *r*^2^ values becoming more variable with increasing contraction intensity was again observed.

**Figure 7.**
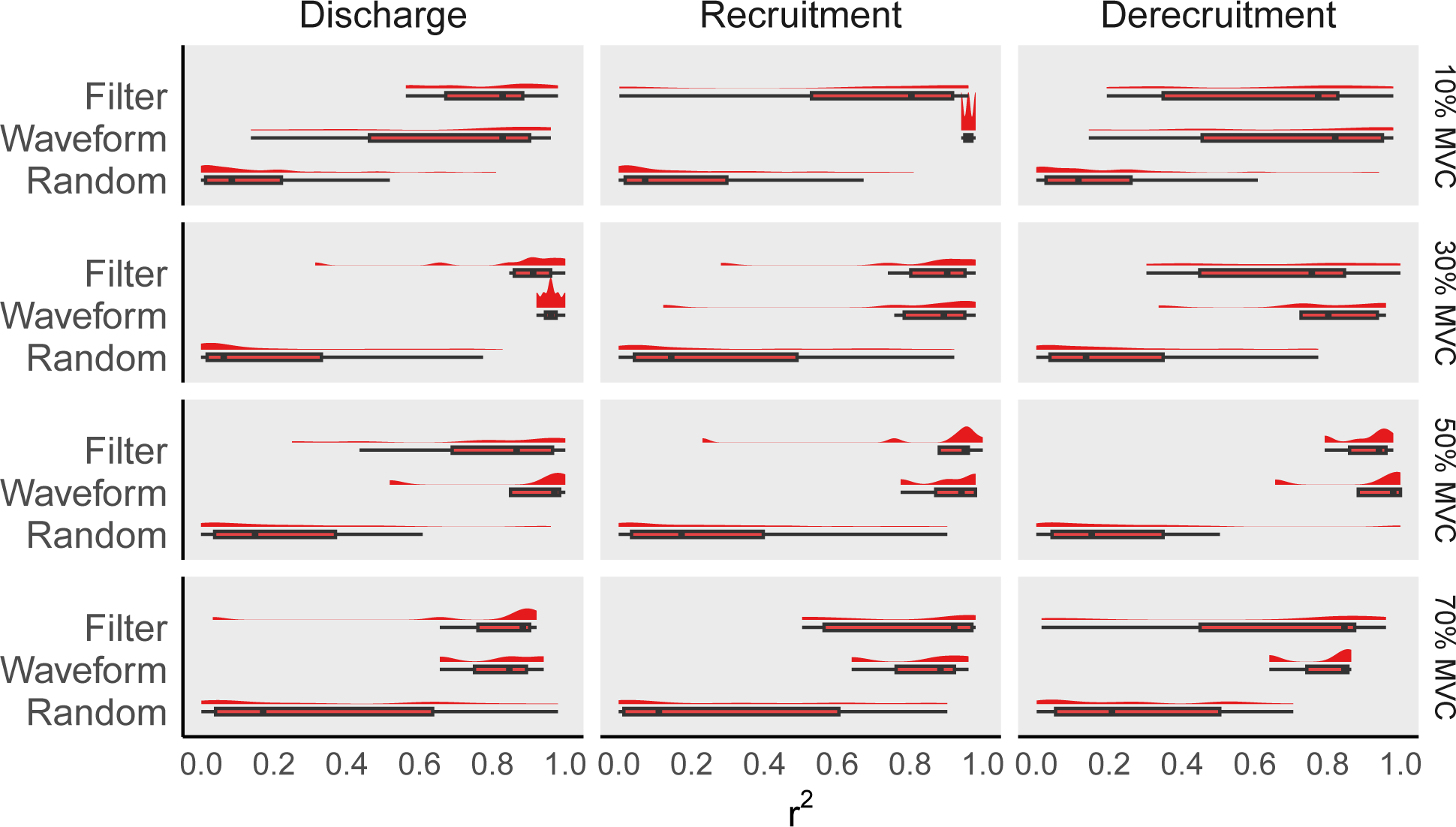
Pearson correlation coefficients for drug data. Box plots and density plots represent the bution of individual subject correlation coefficients for motor units which has been tracked across yproheptadine intervention using the filter, waveform, and random methods. The distribution of r^2^ s is displayed on the abscissa, and distributions are grouped on the ordinate by tracking method. mns represent discharge rate, recruitment threshold and derecruitment threshold, and rows sent the different contraction intensities (10%, 30%, 50% and 70% of MVC).

## DISCUSSION

The purpose of this study was to evaluate the reliability of two common tracking procedures. Intra-session motor unit tracking was performed with the filter tracking method (blind source separation), and inter-session tracking was performed with waveform tracking method (two-dimensional waveform cross correlation). Key findings were: (i) motor unit properties extracted from high intensity contractions demonstrated greater variability than low intensity contractions; (ii) both the filter and waveform methods of tracking substantially outperform multiple iterations of randomly matching motor units which highlights the importance of properly matching motor units across an intervention; (iii) there were no discernible differences in the performance of the filter tracking method and the waveform tracking method where motor units of the tibialis anterior were assessed across the wide range of dorsiflexion intensities; and (iv) although reliability slightly reduced when a pharmacological intervention was employed to reduce discharge rate, there were no discernible differences in the performance of tracking methods with the drug condition. Our results provide evidence that it is possible to track the same motor units in humans with HDsEMG under physiological conditions and following an intervention which reduced the firing rate of motoneurones.

### Stationarity as a principle of motor unit decomposition

Blind source separation algorithms, such as convolution kernel compensation, separate EMG activity into individual motor unit components by detecting the unique motor unit action potential (MUAP) waveform shapes, temporal, and frequency characteristics (7, 15). Once individual MUAP filters are ‘primed’ on these characteristics, they can then be applied to new EMG signal in which the same motor units may be identified. Thus, tracking with the filter technique relies on the stationarity of these characteristics. Similarly, tracking with the waveform technique also requires a level of stationarity in the spatial representation of the identified MUAPs for successful cross correlation (5, 8). Interventions designed to introduce variation in regular physiological function of the central nervous system impact the morphology, temporal, and frequency characteristics of the MUAPs. Therefore, it is of great interest to understand how interventions that alter activity in the motor system, and thus alter MUAPs, affect the reliability of motor unit tracking.

### Similar performance with filter and waveform tracking methods

The filter and waveform tracking techniques performed very similar for the placebo condition. The distribution of change scores for motor unit discharge, motor unit recruitment, and motor unit derecruitment as matched by each technique, was consistently narrow and centred around zero. Measures of reliability, error, and variability were also comparable for both filter and waveform matching techniques. On face value, these reliability findings indicate that one technique is not superior to the other. We also performed iterations of randomly matching motor units together and assessed the reliability of these various random combinations. Compared to filter and waveform based matching of motor units, random matching of units resulted in broader change score distributions and lower correlation coefficients. This highlights the superiority of the filter and waveform techniques for pre-post comparisons over comparisons made between motor units that have been paired at random. A similar analysis has previously been conducted to confirm the accuracy of the waveform tracking between random samples of unmatched motor units from the quadriceps (5), and also from tibialis anterior motor units (8). Collectively, these two studies identified a significant decrease in reliability indices for the randomly matched units when compared to the waveform matched units.

The pharmacological intervention was associated with changes to normal motoneurone activation, which manifested in the reduction of motor unit discharge rate and increasing of derecruitment threshold. Physiologically, it is probable that changes to the intrinsic excitability of spinal motoneurons occurred post-ingestion of the drug (14), which would typically provide an additional source of depolarising current to ionotropic inputs to the motoneuron (23–25). Disruption to 5-HT receptor activity on motoneurones can also reduce firing rate by modulating slow afterhyperpolarisation (sAHP) and medium afterhyperpolarisation (mAHP)(26, 27), which can each contribute changes to temporal, frequency and morphological characteristics of the MUAPs (i.e., stationarity). It is therefore not surprising that the ICCs for the cyproheptadine condition were typically lower than the placebo condition for motor unit discharge rate and derecruitment threshold. Importantly, including an intervention that reduced motor unit discharge did not affect the performance of either filter or waveform tracking techniques in the context of our experiment design. Therefore, the use of either technique would be equally appropriate for matching single motor unit activity with the type of experiment design used in the current study. Furthermore, the ability of the waveform technique to similarly detect pharmacologically induced changes identified via the filter technique adds weight to the growing evidence that 2-D waveform cross-correlation provides a valid and reliable means of longitudinally tracking motor units.

### Motor unit discharge, recruitment, derecruitment, and tracking become more variable with increasing contraction intensity

An increasing number of motor units are recruited to a muscle contraction when the muscle is required to develop greater levels of force (28–31). However, the number of motor units that can be identified by decomposition of HDsEMG typically decreases with increasing contraction intensity (2, 32). Fewer accurately identified motor units can be observed at higher contraction levels, which is likely due to increased superimposition of motor unit action potentials, rather than physiological changes in the number of recruited single motor units (33). Importantly, decomposition of HDsEMG is less successful when trying to identify low threshold motor units during high intensity contractions, causing the sample of successfully decomposed motor units to shift with a bias towards higher threshold units (14). In the present study we observed that the poorest reliability typically occurred at the highest contraction intensity, whereby the greatest variability in tibialis anterior motor unit discharge, recruitment, and derecruitment also occurred at the highest contraction intensity. This finding was consistent across both pharmacological conditions investigated. The source of increasing variability with increasing contraction intensity likely originates from a mixture of low and high threshold motor units being active during higher intensity contractions. (34). Low threshold units require less input to fire and are recruited at lower force levels, while high threshold units require more input to fire and are recruited at higher force levels (29, 30). Therefore, the higher contraction intensities will have the greatest range of motor units active, resulting in a wider range of firing rates and discharge patterns (31, 35). An increase in the variability of motor unit discharge rate with increasing contraction intensity has also previously been observed in vastus lateralis and vastus medialis (4). Overall, as contraction intensity increases, motor unit decomposition and tracking via both techniques becomes more difficult.

### Considerations and future directions

The most direct way by which longitudinal tracking of motor units could be validated would be to concurrently sample HDsEMG and intramuscular EMG signals (22, 36). However, this method has challenges associated with chronic implantation of fine wire electrodes, and the small number of common sources decoded at the surface and into the muscle (36). A pharmacological-based method provides a non-invasive alternative to validate the accuracy and reliability of these methods of tracking motor units. It is important to consider the impact of motor unit yield when discussing the feasibility of longitudinal motor unit tracking. In the current study, 65% of the total yield of tibialis anterior motor units were successfully matched across a 7-day period for the waveform technique. In previous work which tracked tibialis anterior motor units across two sessions via the waveform technique, 30% of the total yield of motor units were successfully tracked over a 4 week period (1). Longitudinal tracking of motor units from the quadriceps muscle group via the waveform method also revealed that the number of successfully matched motor units decreased from ∼45% across two sessions to ∼25% across three sessions (5). Resolving the issue of low yield is critical to aid in the clinical application of tracking motor units over time, whereby disease progression could be reliably monitored in central nervous system conditions that influence motoneurone recruitment, rate coding, and excitability. A further consideration for yield is the muscle from which HDsEMG is being sampled. The current study investigated isometric contractions in tibialis anterior, which is not reflective of all muscles and contraction types. For example, motor unit yield from biceps brachii is typically reported to be far lower than tibialis anterior (2). Therefore, more extensive research is required to characterise the reliability of both upper limb and lower muscles generating a wide range of forces.

## CONCLUSION

This study provides new evidence that it is possible to track the same motor unit in humans with HDsEMG across a wide range of voluntary force levels. Interestingly, there were no discernible differences in the performance of the filter tracking method and the waveform tracking methods under physiological conditions. Although reliability slightly reduced when a pharmacological intervention was employed to reduce discharge rate, there were also no discernible differences in the performance of tracking methods with the drug condition. We observed that the poorest reliability typically occurred at the highest contraction intensity, whereby the greatest variability in tibialis anterior motor unit discharge, recruitment, and derecruitment also occurred at the highest contraction intensity. Collectively, our results provide evidence that it is possible to track the same motor units in humans with HDsEMG under physiological conditions and following an intervention which reduced the firing rate of motoneurones.

